# Pressure overload leads to coronary plaque formation, progression and myocardial events in ApoE^-/-^ mice

**DOI:** 10.1101/562181

**Authors:** Alice Marino, Yi Zhang, Luisa Rubinelli, James E Ip, Annarita Di Lorenzo

## Abstract

Hypercholesterolemia and hypertension are two major risk factors for coronary artery diseases, which remain the major cause of mortality in the industrialized world. Current animal models of atherosclerosis do not recapitulate coronary plaque disruption, thrombosis and myocardial infarction occurring in human. Recently, we demonstrated that exposure of the heart to high pressure, by transverse aortic constriction (TAC), induced coronary lesions in ApoE^-/-^ mice on chow diet.

The aim of this study was to characterize the magnitude and location of coronary lesions in ApoE^-/-^ mice post-TAC, and to assess the susceptibility of coronary plaque to disruption, leading to myocardial events.

Here we describe a reliable pathological condition in mice characterized by the development of coronary lesions and its progression leading to myocardial infarction, which better recapitulate human disease. Following TAC surgery, about 90% of ApoE^-/-^ mice develop coronary lesions, especially in the left anterior descending artery, with 59% of the mice manifesting different magnitude of LAD stenosis. Myocardial events, identified in 74% of the mice, were mainly due to coronary plaque thrombosis and occlusion.

That TAC induces the development and progression of coronary lesions in ApoE^-/-^ mice leading to myocardial events represents a novel and important tool to investigate the development of coronary lesions and its sequelae in a setting that better resemble human conditions.

## Introduction

Coronary artery disease (CAD) remains the leading cause of morbidity and mortality in the US, accounting for more than 40% of deaths due to cardiovascular disease. Statins have been partially successful, suggesting the need for alternative therapeutic options(1). In this regard, the development of novel therapeutic approaches is impeded, in part, by the use of animal models of atherosclerosis that do not fully mirror the features of human CAD(2).

Since 1994, ApoE^-/-^ mice(3) have been widely used to study the pathogenesis of atherosclerosis, and to develop pharmacologic and other interventional therapeutic approaches(4-6). In spite of high plasma cholesterol(6), ApoE^-/-^ mice develop minimal atherosclerosis on a regular diet(3). Thus, to induce atherosclerosis, ApoE^-/-^ mice, have been fed with Western-type diet(3), resulting in plasma cholesterol levels ≥1400 mg/dl, which is markedly higher than those reported in humans(7). Subsequently, the same approach was used for LDL receptor knockout (LDLR^-/-^) mice(8). ApoE^-/-^ and LDLR^-/-^ mice fed high fat diets develop extensive lesions in the aortic root, the aorta and its principal branches; however they fail to develop atherosclerosis in the coronary arteries, a major affected site in humans(9). Moreover, in these models coronary plaque progression and its sequelae are not observed.

To overcome this drawback and induce coronary lesions, genetic mouse models were generated by excising srb1 gene (scavenger receptor class B type I, SR-BI)(10), ldlr gene (LDLR)(11) on ApoE^-/-^ background. However, long-term high fat diet was still required to induce coronary atherosclerosis in the LDLR^-/-^/ApoE^-/-^ mice, and remained not sufficient to elicit myocardial infarction (MI). Indeed, additional hypoxic or mental stress was necessary to cause MI, which occurred in only 40% of the mice(11). SRBI^-/-^ - ApoE^-/-^ mice developed coronary lesions on regular diet, but they had an incredible short lifetime (8 weeks) due to a complex pathological condition making this model not amenable to pre-clinical studies(10). Another attempt to induce coronary lesions in mice was made by heterozygous mutation of fibrillin-1 gene on ApoE^−/−^ background(12). Although coronary lesions formed, stroke was the primary cause of death in more than 60% of the mice(12). Thus, the clinically relevant events of coronary plaque rupture and thrombosis cannot be systematically studied in existing mouse models.

Established risk factors for CAD are hypercholesterolemia and high blood pressure(13, 14). In line with clinical findings, transverse aortic constriction (TAC) in ApoE^-/-^ mice, on regular chow diet, induced the development of atherosclerotic lesions in the vascular segments exposed to high pressure, including right carotid artery and coronary arteries, but not in the vessels at low pressure (i.e., left carotid artery and descending aorta)(15). The aim of this study was to define the magnitude and location of coronary lesions in ApoE^-/-^ mice post-TAC, and to investigate whether the susceptibility of coronary plaque to disruption was reproducible and accompanied by “human-like” events, such as myocardial infarction and/or sudden death.

Here, we report that TAC-induced coronary atherosclerosis can progress to plaque thrombosis, occlusion and MI. At 8-week post-TAC, more than 90% of ApoE^-/-^ mice developed atherosclerotic lesions in the coronary arteries, particularly in the left anterior descending artery (LAD). About 60% of the mice presented LAD stenosis.

When subjected to physical stress conditions (i.e. treadmill), more than 70% of the TAC-operated ApoE^-/-^ mice died and the majority showed striking evidence of previous myocardial infarction. Systematic and robust histological analysis revealed that MI was caused mainly by coronary occlusion or plaque rupture and/or erosion.

Our study demonstrates, for the first time, that ApoE^-/-^ mice are susceptible to coronary lesion formation and its sequela if chronically exposed to elevated cholesterol (∼350 mg/ml) and increased pressure afterload. The formation of coronary lesions in ApoE^-/-^ mice following pressure overload makes them of great experimental value. The combined phenotype (high cholesterol/pressure overload with coronary lesions) provides a baseline against which either detrimental or protective genetic and environmental factors, as well as drugs targeting atherosclerosis, can be investigated.

## Results

### Characterization and distribution of coronary lesions in ApoE^-/-^ mice at 8 weeks post-TAC

TAC surgery was performed on ApoE^-/-^ mice followed by longitudinal assessment of different parameters, including magnitude and location of coronary lesions, electrocardiograms (ECG), echocardiography and histology of the hearts (Fig. 1A). At 8-weeks post-TAC, histological analysis of heart sections from the aortic valve (AV) to the apex was performed to identify the presence of coronary lesions, the degree of stenosis and the lipid content of the plaques (Fig. 1B-E). The majority of early LAD lesions and its branches (∼80%) presented lipid deposition in both intima and media (Fig. 1B and Supplementary Figure 1A). Macrophages with different degrees of lipid accumulation were observed in the intima and in the media (Fig. 1B, early lesions, arrow and arrowhead respectively). In 20% of the early lesions, the accumulation of the lipids was limited to the intima or to the media (Supplementary Figure 1A). Just as endothelial cells (EC), activated smooth muscle cells (SMC) can express adhesion molecules, such as intercellular adhesion molecule (ICAM) and vascular cell adhesion molecule (VCAM), and therefore can interact and retain monocytes /macrophages into the atherosclerotic artery(16). Interestingly, the initial accumulation of lipids into the coronary wall seems to be different throughout the coronary tree. Specifically, the media of smaller coronary arteries in the apex of the heart (i.e. distal LAD, marginal and diagonal arteries) presented greater lipid accrual compared to the proximal LAD (Supplementary Figure 1B), suggesting a different role of SMC in distinct coronary artery segments during the initial phase of atherosclerosis. Altogether, these findings suggest similar aspects in the onset of coronary atherosclerosis in mice and human, including the initial local accumulation of lipids in the intima and media, neointimal thickening, and recruitment of macrophages, which later develop into foam cells. The majority of advanced coronary lesions presented a significant neointimal thickening with lipid accumulation, and different degree of macrophage infiltration. Fig. 1B shows two examples of advanced lesions, one causing <50% of luminal stenosis, with pronounced neointimal thickening and infiltrated macrophages (advanced lesion 1); and another presenting >50% of stenosis (advanced lesion 2), with robust macrophage infiltration forming foam cells into the intima (asterisks) and few in the media (arrowhead).

**Figure 1.**
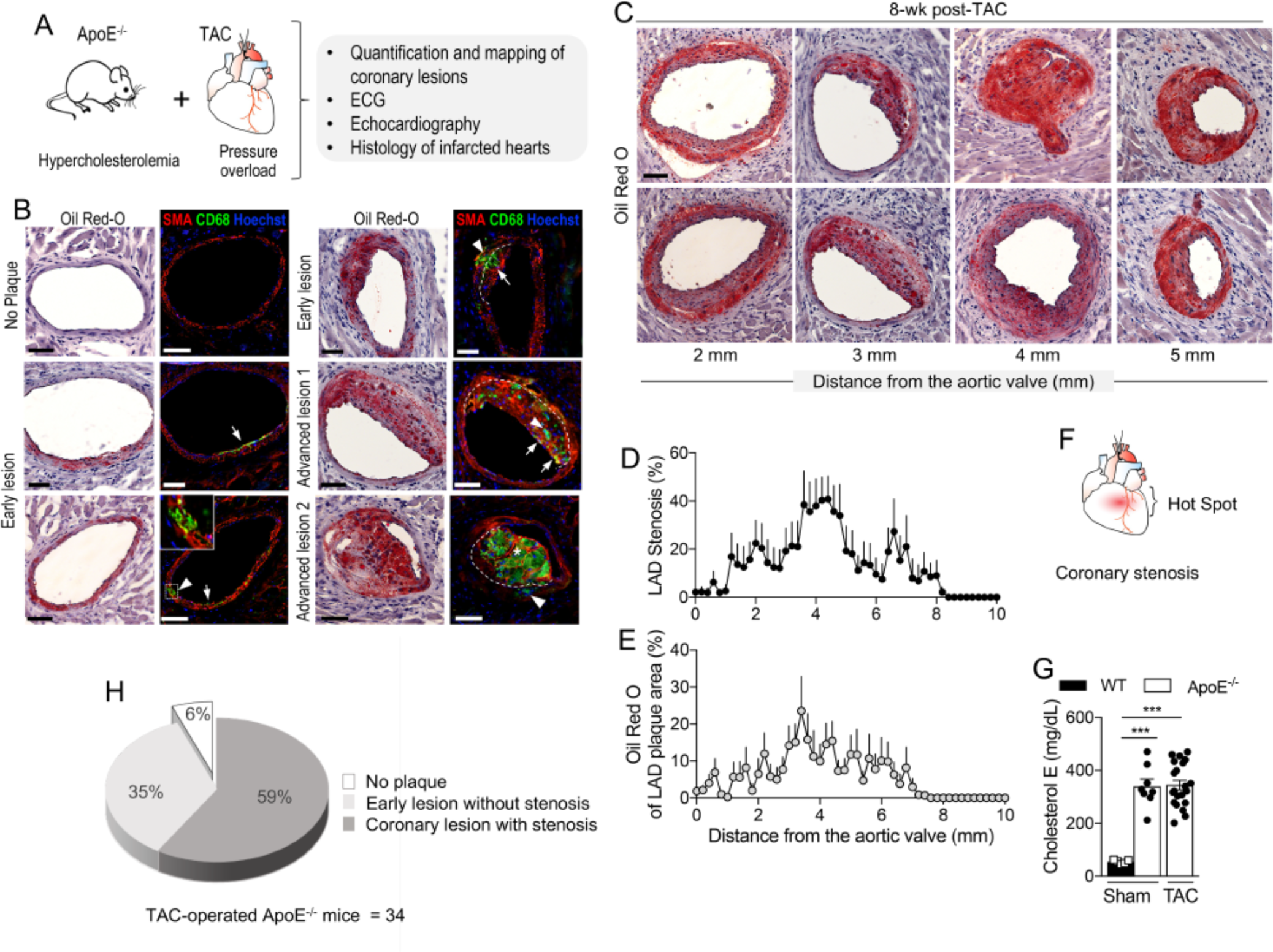
Characterization and distribution of coronary lesions in ApoE^-/-^ mice post-TAC. (A) Scheme of the experimental design. Transverse aortic constriction (TAC) was performed in male ApoE^-/-^ mice (∼25g of body weight) on chow diet, followed by histological, echocardiographic and electrophysiological analysis at different time points post-TAC. (B) At 8-wk post-TAC, sequential myocardial sections were stained with oil red-O (binding to lipids) & hematoxylin, and immunofluorescently stained with α-SMA (SMC), CD68 (monocyte/macrophage) and Hoechst (nuclei). Oil red-O evidenced lipid accumulation in the vascular wall, including endothelial cells (EC), SMC, and macrophages, of early and advanced coronary lesions. Consecutive heart sections immunofluorescently labeled showed macrophage (green) infiltration into the intima (white arrows) and media (white arrowheads) of the coronary arteries. (C) Representative images of the coronary plaques in the LAD of 8-wk TAC-operated ApoE^-/-^ mice at different distance from the aortic valve (AV) showed different degree of stenosis. (D) Quantification of LAD stenosis from the AV to the apex of 8-wk TAC-operated ApoE^-/-^ mice (n=10). (E) Base-to-apex lipid accumulation into the same LAD lesions, expressed as percentage of the plaque area (n=10). (F) Cartoon of the “hot spot” of occurrence of coronary plaques. (G) Total plasma cholesterol levels of WT (n=5), sham- and TAC ApoE^-/-^ mice (n=8 and 20, respectively; plasma was collected between 1 and 11 weeks post-surgery). (H) The pie chart shows the percentage of ApoE^-/-^ mice with no lesions, early lesions with no stenosis, and coronary lesions with stenosis (n=34). ***, P<0.001, One-way ANOVA.

Next, 8-weeks post-TAC, we quantified the extent of LAD stenosis in sequential myocardial sections from the AV to the apex (Fig. 1C-D). The highest stenosis was observed along the proximal coronary vasculature, specifically within 5 mm from the AV (Fig. 1D). The lipid plaque composition followed a similar longitudinal profile (Fig. 1E). Based on the analysis of the LAD stenosis, we defined the proximal area of the LAD as “hot spot” (Fig. 1F). It is noteworthy to mention that the “hot spot” in this murine model of coronary lesions is similar to the one observed in human coronary arteries(17). As expected, cholesterol levels in ApoE^-/-^ mice were significantly higher than WT mice(6), and were not different between sham- and TAC-operated ApoE^-/-^ mice (Fig. 1G), suggesting that the high pressure following aortic banding, in combination with moderate high cholesterol (∼350 mg/dL), is responsible for coronary plaque formation. The analysis of coronary lesions in the heart of 34 TAC-operated ApoE^-/-^ mice revealed that 59% developed LAD stenosis, 35% showed early lesions without stenosis, and only 6% had no evidence of atherosclerosis (Fig. 1H).

### Evidence of myocardial infarction in ApoE^-/-^ mice post-TAC exposed to physical stress

By 10-weeks post-TAC, more than 60% of ApoE^-/-^ mice had died. Post-mortem examination of some of these mice revealed areas of pale myocardium, suggestive of MI.

Next, ApoE^-/-^ mice were physically stressed by treadmill run (once per week), starting at 8 weeks post-TAC, when coronary lesions were established. During this protocol, echocardiographic analysis and ECG telemetry were performed (Fig. 2A). After 25 weeks, 17% of the TAC-operated ApoE^-/-^ mice survived, 17% died before the physical stress protocol, and 66% died during the physical stress protocol (Fig. 2B). The Kaplan Meier curve shows that ∼50% of the ApoE^-/-^ mice died by week 14, and about 80% of the mice by week 25 (Fig. 2C). Post-mortem analysis of these mice also revealed pale myocardial areas suggestive of infarct (Fig. 2D).

**Figure 2.**
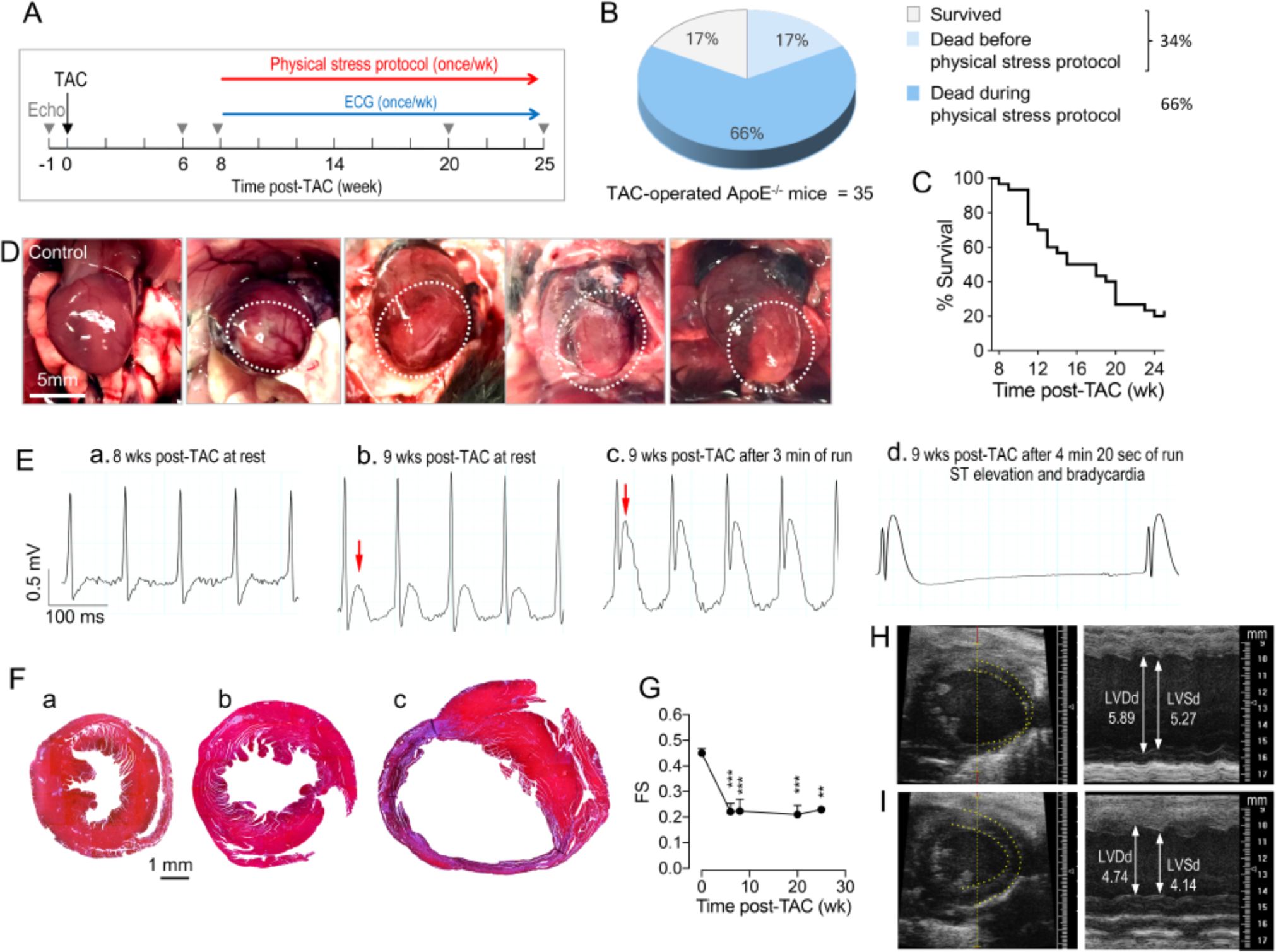
Physical stress induces myocardial events in ApoE^-/-^ mice post-TAC. (A) Scheme of the physical stress-induced MI protocol: starting at 8 weeks post-TAC, once per week ApoE^-/-^ mice were physically stressed by a treadmill run (4 min, at 13 m/min). ECG traces were recorded at baseline, during and 48h after the run on the treadmill by using radiotelemetry devices. Echocardiographic analysis was performed at baseline and at 6, 8, 20 and 25 weeks post-TAC, as indicated by the gray arrowheads. (B) Percentage of mice that died before, during the stress test protocol, and survived between 8 and 25 weeks post-TAC (n=35). (C) Survival curve of ApoE^-/-^ mice post-TAC dead during the physical stress-induced MI protocol (from 8 to 25 weeks post-TAC; n=29). (D) Images of the infarcted ApoE^-/-^ hearts at post-mortem examination. Dotted circles indicate the pale myocardium, index of infarct. (E) Representative ECG tracing o TAC-operated ApoE^-/-^ subjected to the physical stress protocol: (a) during rest; (b) the same mouse a week later shows ST elevation (red arrow) at rest; (c) following treadmill run ST-elevation was further increased. (d) The mouse died with remarkable signs of ST elevation and profound bradycardia. (F) Masson’s trichrome staining of (a) banded WT heart at 12-week post-TAC (b) TAC-operated ApoE^-/-^ mouse survived the physical stress (25-week post-TAC) (c) TAC-operated ApoE^-/-^ mouse died during physical stress at 12-week post-TAC. (G) Fractional shortening (FS) measured at the indicated time points after TAC. (n≥5) **P < 0.01, ***P < 0.001, one way-ANOVA. Representative images of two-dimensional M-mode echocardiography of the LV of TAC-operated ApoE^-/-^ mice (H) without infarct and (I) with infarct, showing the thin ventricular wall in the apex (yellow dotted lines).

Eight TAC-operated ApoE^-/-^ mice were implanted with radiotelemetry devices to measure single lead ECG once per week, at rest, during and 48h after physical exercise. ECG traces showed a clear ST-elevation in 3 mice, and arrhythmic events in the remaining. The latter included accelerated idioventricular rhythm, which is associated with myocardial ischemia, and sudden profound bradycardia (sinus bradycardia and atrioventricular block). In the ECG depicted in Figure 2E, the ApoE^-/-^ mouse at 9-weeks post-TAC showed ST elevation at rest (Fig. 2Eb), that became more prominent during exercise (Fig. 2Ec), and the mouse died immediately after running with associated sudden bradycardia (Fig. 2Ed).

Masson’s trichrome staining of banded hearts from mice that died with ST-elevations revealed an extended cardiac fibrosis due to infarction damage. As controls, the Masson’s trichrome staining was performed on the banded hearts of WT (Fig. 2Fa) and survived ApoE^-/-^ (Fig. 2Fb) mice at the same time point post-TAC showing some degree of fibrosis, as expected in pressure-overloaded hearts, but significantly less than the infarcted ApoE^-/-^ hearts (Fig. 2Fc).

Lastly, as expected fractional shortening was decreased following TAC (8 to 25 weeks, Fig. 2G). Echocardiographic images showed left ventricular dilation, reduced function and evident ventricular wall thinning in infarcted hearts (Fig. 2H) compared to non-infarcted hearts (Fig. 2I) of TAC-operated ApoE^-/-^ mice.

### Evidence of plaque disruption, thrombosis and MI TAC-operated ApoE^-/-^ mice

Hearts of TAC-operated ApoE^-/-^ mice that died during the physical stress protocol were systematically sectioned from the AV to the apex and stained with hematoxylin and eosin (H&E). This first histological examination allowed us to identify points of plaque occlusion or disruption for further analysis. H&E staining of sequential myocardial sections revealed two severe atherosclerotic lesions in the LAD at 2.95 mm from the AV (Fig. 3A, panel 4, asterisks). Juxtaposed to the right plaque, it is noticeable a “white” thrombus (inset i, arrowheads) and red blood cells confined within the narrow lumen. Next, we sought to determine whether plaque rupture and/or erosion caused the thrombotic event. Thus, myocardial sections, upstream and downstream to the one in panel 4 (Fig. 3A), were stained for CD41, to identify platelets aggregates, alpha-smooth muscle actin (α-SMA) for SMC, and collagen I (Col I, marker of extracellular matrix) to define the fibrous cap of the plaque (Fig. 3C). The immunofluorescent staining of the LAD showed a thin but continuous fibrous cap of the left plaque (inset i), and two points of rupture in the fibrous cap of the right plaque, evidenced by the interrupted Col I staining (Fig. 3C, inset ii). In Fig. 3D, CD41 staining (red) of a consecutive section, confirmed the presence of a thrombus, which extended from the lumen to inside the ruptured plaque (Fig. 3Dii). These events are also depicted in the cartoon in Fig. 3E. It appears also clear that the thrombus is confined to the lumen in front of the left plaque (panel 3D, F and relative insets i), with thin and undisrupted fibrous cap. Perhaps erosion might have occurred as the thrombus juxtaposes to the luminal surface of the left plaque.

**Figure 3.**
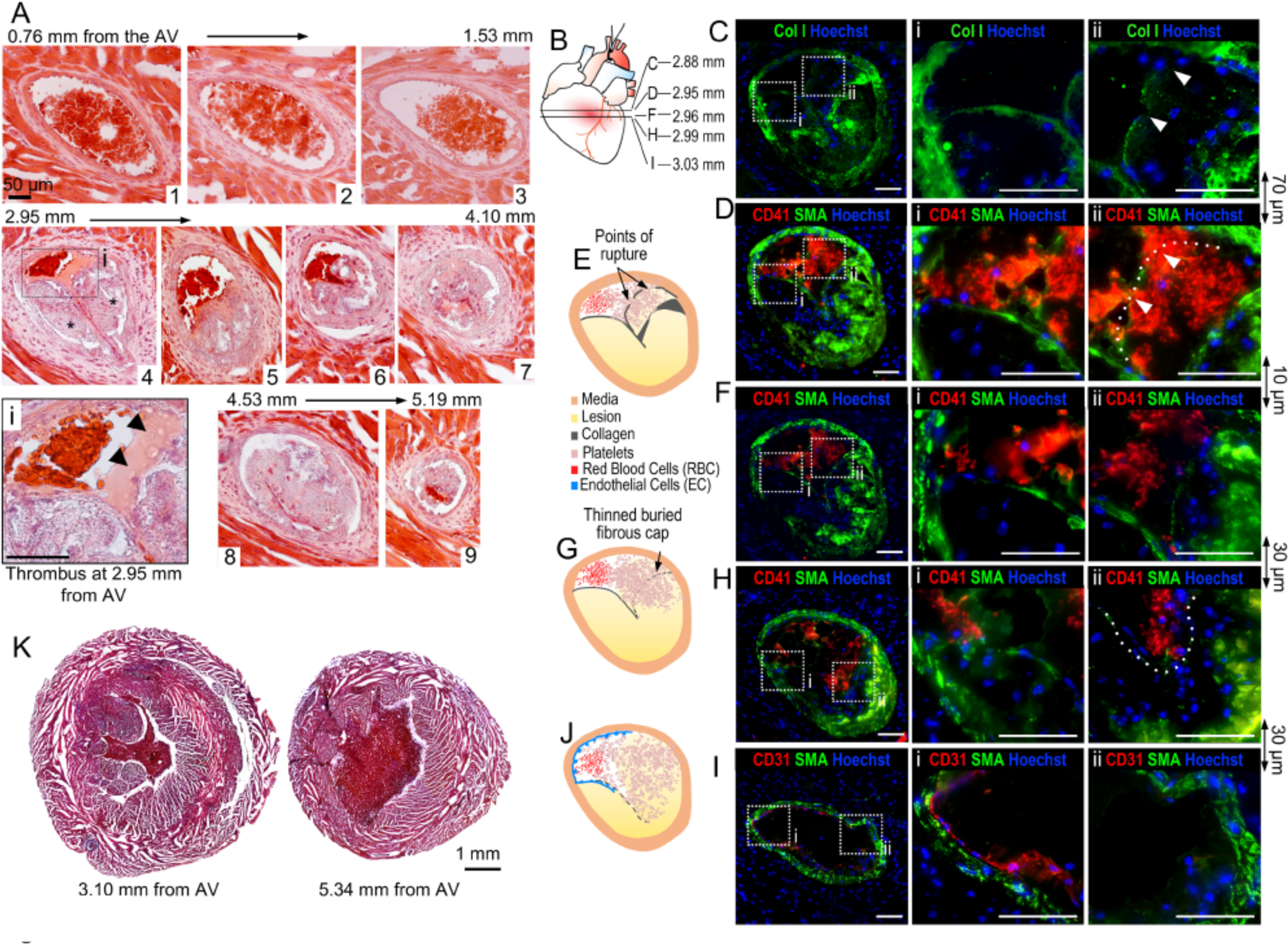
Coronary plaque rupture and thrombosis in the LAD of TAC-operated ApoE^-/-^ mice. Sequential myocardial histology of TAC operated ApoE^-/-^ mice died during physical stress protocol. (A) H&E staining of myocardial sections showing the LAD. At 2.95 mm from the AV there are two major lesions and a thrombus (light pink) juxtaposed to the plaque on the right (asterisks, panel 4 and inset i). (i) Magnified image of the thrombus with visible platelets (arrowheads), and red blood cells confined in a narrow lumen space. (B) Scheme of the localization of immunofluorescently stained myocardial sections proximal and distal to the thrombus shown in A, panel 4. (C) IF staining of the LAD for collagen type-I (Col I, green) and Hoechst (nuclei, blue), showing a thin and continuous fibrous cap on the left plaque (i) and a thin and discontinuous fibrous cap of the right plaque (ii). The two ruptured points are indicated by two arrowheads. (D) IF staining for CD41 (platelets, red), αSMA (smooth muscle cells, green) and Hoechst (blue) shows luminal thrombus juxtaposed to the left plaque (inset i). In the inset (ii) the thrombus is present in front of and within the right-ruptured plaque (arrowheads and dotted line). (E) Schematic representation of the IF staining in panels D and F of the ruptured plaque and thrombus in the LAD.(F) CD41, αSMA and Hoechst IF staining proximal consecutive section of the LAD showing the same pattern of thrombus distribution in the lumen and within the ruptured plaque. (G) Schematic representation of the IF staining in panels H of the ruptured plaque and thrombus in the LAD. (H) Consecutive distal sections showing platelet infiltration and aggregation deeper into the plaque. Part of the right plaque detached from the arterial wall (dotted line). (I) IF staining for CD31 (EC) and αSMA of a consecutive distal section shows CD31 positive staining only in the upper left corner of the LAD, whereas is absent in the remaining LAD, where the ruptured plaque material and clots broke away from the wall. (J) Schematic representation of the IF staining in panels I of the disrupted endothelium in the LAD. In all the panels the scale bars represents 50 µm. (K) Masson’s trichrome staining of the medium and apex of the heart at the indicated distance from the AV.

In distal myocardial sections, the thrombus further extended into the ruptured plaque in intimate contact with the thrombogenic material (Fig. 3H-I and relative cartoons G and J). Moreover, CD31 staining (endothelial cells) showed the lack of the endothelium on the right-ruptured plaque, while a positive staining can be observed in the upper-left part of the lumen (Fig. 3I). Masson’s trichrome staining of cross sections from in the medium and apex of the heart revealed some perivascular and interstitial fibrosis but no signs of chronic MI(Fig. 3K).

Another example of coronary plaque rupture is shown in Supplemental figure 2. It is noteworthy to mention that platelet aggregation it is not present in proximal or distal myocardial sections to 2.95 mm from AV, suggesting that the observed thrombus did not result from stagnant blood flow in the coronary vasculature. Masson’s trichrome demonstrated minimal cardiac fibrosis suggesting the onset of an acute MI due to plaque rupture (Supplementary Fig. 2P).

A significant cause of acute coronary syndrome is superficial plaque erosion (18). This event involves fibrous, rather than lipid-rich plaque, and disturbed flow may contribute to endothelial pro-thrombotic phenotype and apoptosis (19, 20). H&E staining of sequential myocardial sections of a TAC-operated ApoE^-/-^ mouse that died during the physical stress protocol showed the presence of “white” thrombus at about 3 mm from the AV (Fig. 4A, B). Immunofluorescent staining of a consecutive myocardial sections for CD41 corroborated this observation. Whereas at 2.57 mm from the AV -upstream the point of erosion-no thrombus was observed (Fig. 4C and D), at about 3 mm from AV, CD41 staining evidenced the presence of thrombus in the lumen, adjacent to, but not inside the plaque (Fig. 4F and G). This observation was supported by Col I staining showing a thick fibrous cap, thus excluding the rupture of the plaque (Figure 4H and I). The lack of notable endothelial cell nuclei in Figure 4H and I suggested erosion of the endothelium. CD31 staining confirmed the absence of the endothelium on the plaque in contact with the thrombus (Fig. 4Jii, arrowheads), whereas a positive CD31 staining was present on the opposite site of the LAD (Fig. 4Ji, arrows), where the thrombus was absent. These data suggest that coronary plaque erosion triggered the formation of thrombus; contributing to MI. Masson’s trichrome staining of the heart with minimal fibrosis suggests the onset of an acute MI due to erosion rather than chronic infarct (Fig. 4L).

**Figure 4.**
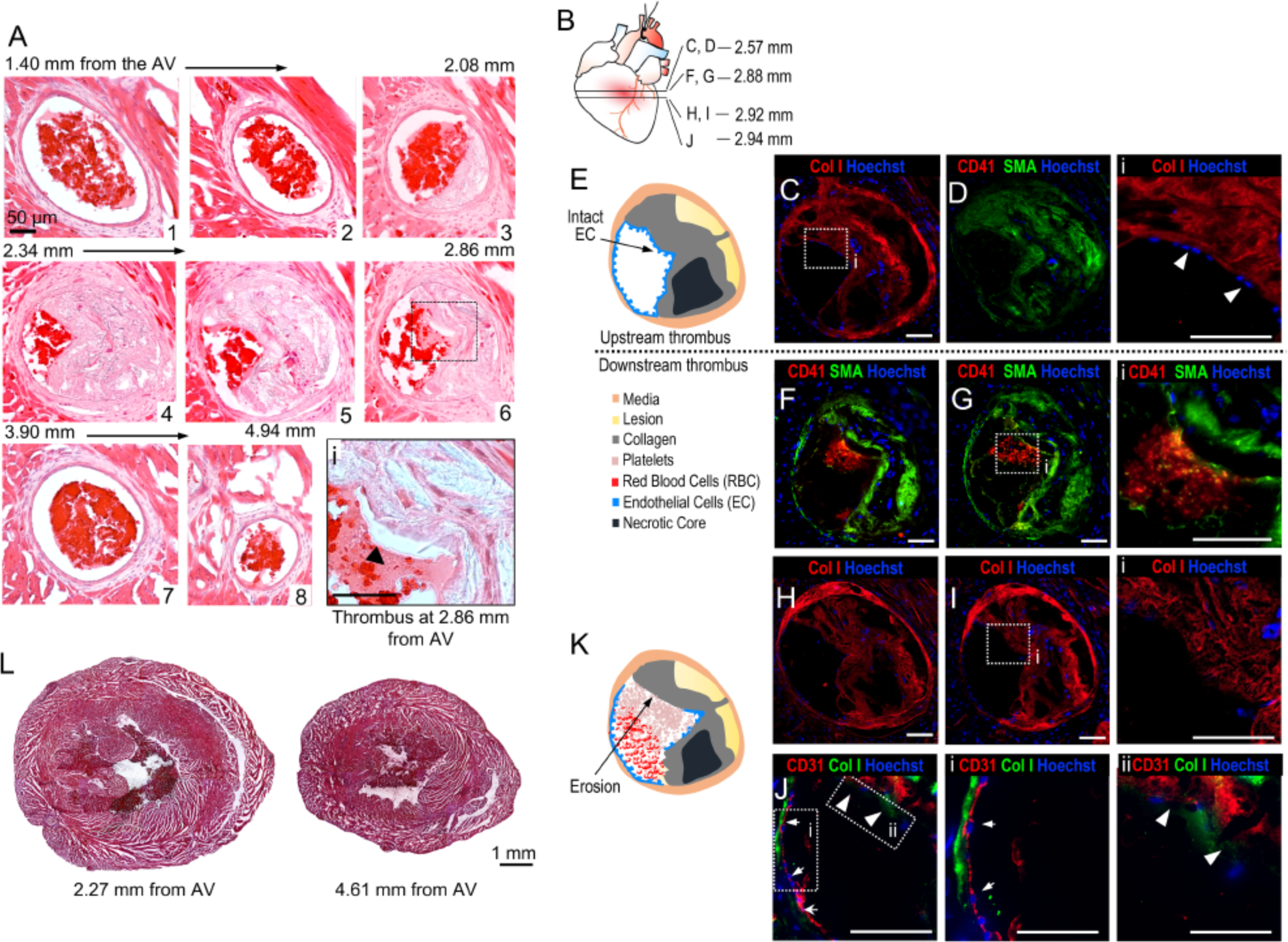
Coronary plaque erosion and thrombosis in ApoE^-/-^ mice post-TAC. (A) H&E staining of sequential myocardial sections of ApoE^-/-^ mouse died during the physical stress protocol. At 2.86 mm from AV (panel 6) is noticeable the thrombus (light pink) in contact with the coronary plaque of the LAD. The magnification of the thrombus is shown in panel i where the arrowheads indicate visible platelets. (B) Scheme of the localization of immunofluorescently labeled heart sections proximal and distal to the section at 2.86mm from the AV. (C, D) IF staining of proximal section to erosion for Col I, CD41, αSMA and Hoechst showed the absence of platelet aggregation. Hoechst staining shows the presence of elongated endothelial cell nuclei (C, inset i). (E) Schematic representation of the IF staining upstream the erosion shown in C and D. (F-I) IF for Col I, CD41, αSMA and Hoechst of myocardial sections displaying the thrombus in the lumen, adjacent to but not inside the plaque (panel G, inset i). The fibrous cap (Col I and αSMA staining) adjacent to the thrombus is very thick and lacks of endothelial cell nuclei (panel I, inset i). (J) The staining for CD31 shows the presence of endothelial cells on the left side of the LAD free from platelet aggregates (panel J, inset i, arrows) but the absence of endothelial cells on the right side of the plaque in contact with the thrombus (panel J, inset ii, arrowheads). (K) Schematic representation of the IF staining of the eroded plaque with the thrombus in the LAD shown in H and J. Scale bars: 50 µm. (L) Masson’s trichrome staining of the medium and apex of the heart at the indicated distance from the AV.

### Evidence of MI caused by coronary occlusion in TAC-operated ApoE^-/-^ mice

Histological analysis of the heart showed coronary stenosis atherosclerosis and no evidence of plaque disruption and thrombosis (Fig. 5A). Specifically, it was evident the occlusion of LAD, and/or its branches (i.e. diagonal and marginal arteries). H&E, Masson’s trichrome and Oil Red-O staining of myocardial sections from the base to apex revealed the occlusion of the distal LAD and diagonal arteries (Fig. 5B-D). These advanced plaques were filled with cholesterol crystals (Fig. 5C, insets i and ii) and lipids (Fig. 5D). Five of the six –TAC-operated ApoE^-/-^ mice presenting with coronary occlusions were also evaluated with ECG telemetry during the physical stress protocol. Two mice showed ST-elevation and three mice manifested an accelerated idioventricular rhythm, suggestive of myocardial ischemia.

**Figure 5.**
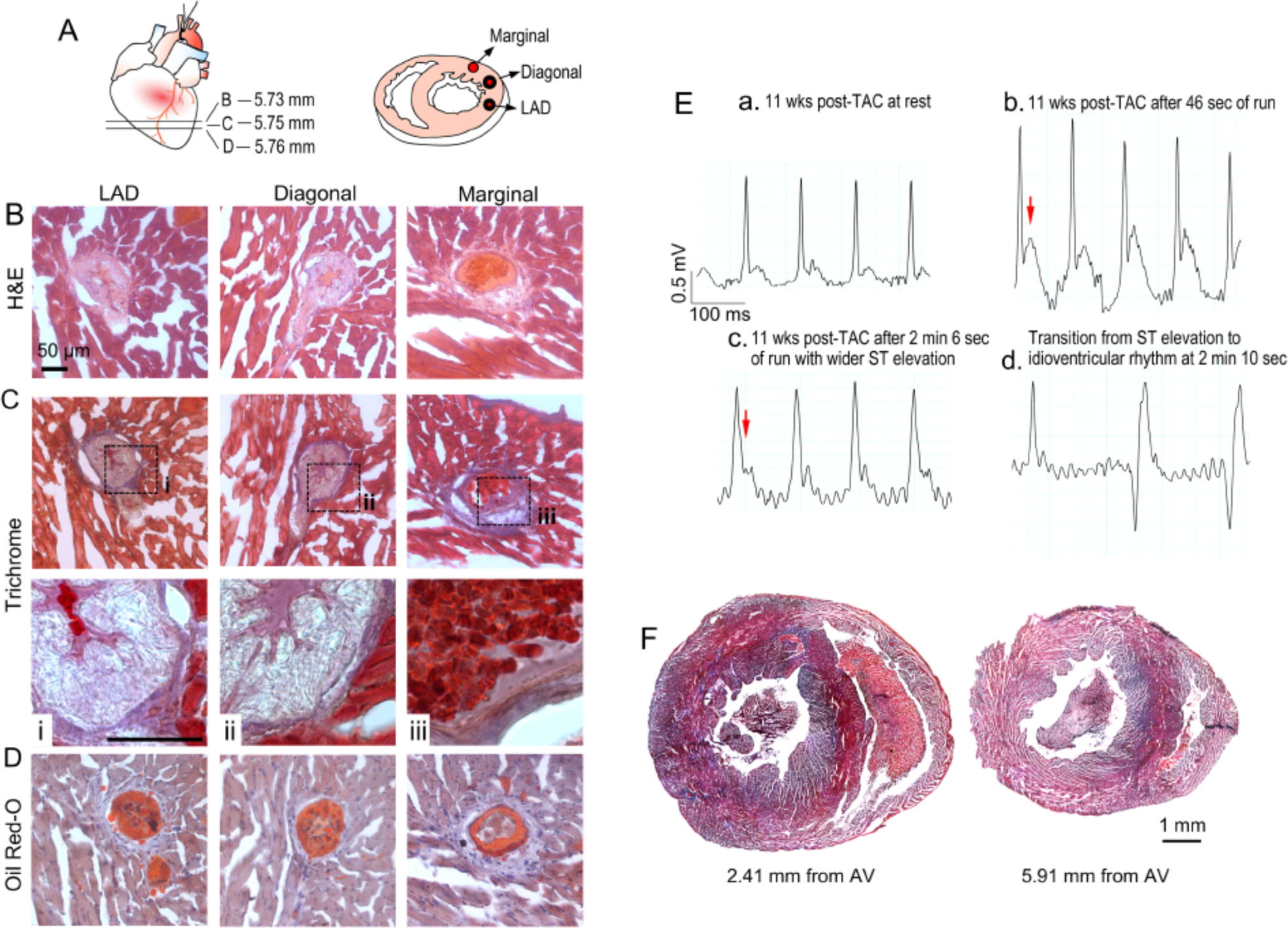
Coronary occlusions without thrombus in epicardial coronary arteries of ApoE^-/-^ mice post-TAC affected by MI. (A) Scheme of the localization of heart sections stained with H&E, Masson’s Trichrome, and Oil-red-O in the apex, and cartoon (right) of the occluded coronary arteries. (B) H&E, (C and insets i, ii, iii) Masson trichrome, and (D) oil red-O staining of distal coronary arteries showing remarkable atherosclerotic occlusion, and likely vasospasm, of the LAD and diagonal arteries, but not of the marginal artery, at 5.73 mm from AV. (E, a) ECG tracing measured at 11 weeks post-TAC in ApoE^-/-^ mouse at rest. (b) ST elevation occurred after 46 seconds of run and (c) it became wider after 2 min of run. (d) The mouse died after a transition from sinus tachycardia with ST elevation to an idioventricular rhythm. Scale bars: 50 µm. (F) Masson’s trichrome staining of the medium and apex of the heart at the indicated distance from the AV.

Figure 5E shows the ECG changes of an ApoE^-/-^ mouse at 11 weeks post-TAC (a) at rest and (b and c) during the run on the treadmill. ST elevation was evident already at 46 sec of run (Fig. 5E,b). After 2 minutes of running, the QRS complex became wider (c), finally leading to an accelerated idioventricular rhythm (d) and death. In addition to the coronary occlusion, coronary spasm could also have contributed to the myocardial event.

Interestingly, Masson’s trichrome staining of the hearts of all six mice that died during the physical stress protocol and presenting with coronary occlusion showed evidence of extended fibrosis (Fig. 5F), suggesting chronic myocardial infarctions, which might have contributed to stress-induced death from ischemic cardiomyopathy.

### Evidence of coronary embolism in TAC-operated ApoE^-/-^ mice

Coronary artery embolism is a rare non-atherosclerotic cause of acute MI, attributed mainly to atrial fibrillation, following by dilated cardiomyopathy and valvular heart disease(21). In most cases, coronary artery embolism involves the distal LAD, causing transmural MI(22). Interestingly, we found a few cases of abrupt occlusion of coronary arteries without significant atherosclerosis. H&E staining of myocardial sections from the base to the apex clearly showed the absence of atherosclerotic plaques, with abrupt interruption of blood flow in the distal LAD, at approximately 6 mm from the AV (Fig. 6A). It is possible to notice red blood cells in panel 7, and the thrombotic occlusion in the distal panel 8 of Fig. 6A. CD41 immunofluorescent staining displayed platelet aggregation occluding the entire lumen and disrupting the continuity of the endothelium in some areas (Fig. 6F-I) but not upstream the thrombus (Fig. 6C-E). In Fig. 6J Masson’s trichrome staining revealed endocardial fibrosis and left ventricular dilation, which may have contributed to the formation of thrombosis that caused the coronary embolism. Because of the absence of significant atherosclerosis in this mouse and the presence of thrombus creating an abrupt occlusion of the distal LAD, we classified this case as coronary embolism.

**Figure 6.**
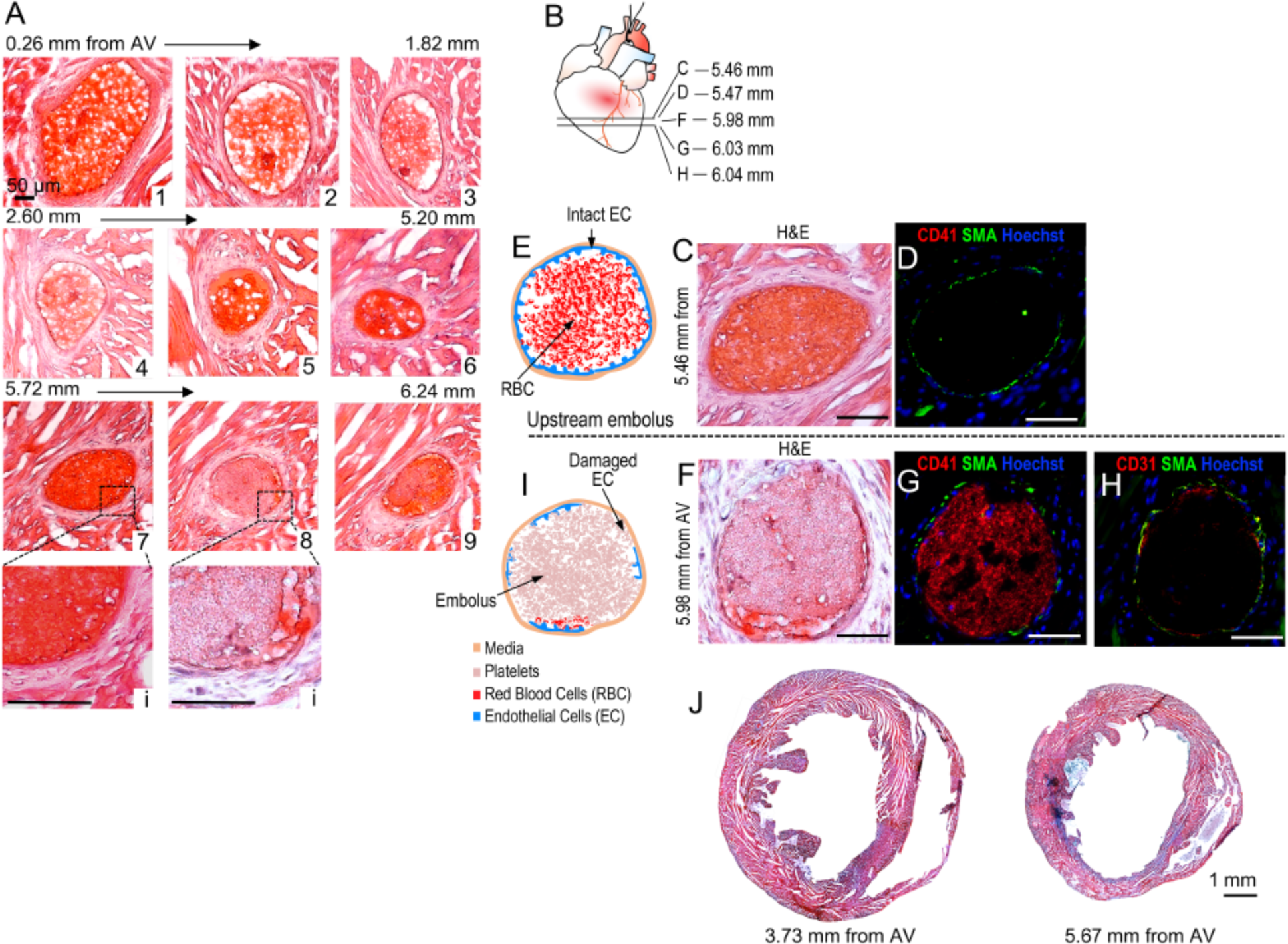
Coronary artery embolism in the distal LAD of TAC-operated ApoE^-/-^ mice. (A) H&E staining of myocardial sections of ApoE^-/-^ mouse died during the physical stress protocol. (B) Scheme of the localization of immunofluorescently labeled heart sections proximal and distal to the section with thrombus at 5.98 mm from the AV, shown in A, panel 8. (C,D) H&E and IF staining of consecutive myocardial sections upstream the embolus. CD41 staining is negative confirming the absence of thrombus at about 0.5 mm upstream the myocardial section in A, panel 8. (E) Schematic representation of the LAD staining in panels C and D. (F-H) Embolus was detected by H&E and IF staining for Col I, CD41, CD31, αSMA and Hoechst. The endothelium in the correspondence of the embolus appears partially damaged, most likely by the thrombus since there are no evident lesions. (I) Scheme of the immunofluorescently labeled LAD shown in F-H. Scale bars: 50 µm. (J) Masson’s trichrome staining of the medium and apex of the heart at the indicated distance from the AV.

### Causes of MI in TAC-operated ApoE^-/-^ mice subjected to the physical stress protocol

Based on systematic histological analysis, combined with ECG and echocardiographic measurements, we found that 74% of TAC-operated ApoE^-/-^ mice died during the physical stress protocol showed evidence of MI (Fig. 7A). In 17% of the cases we were not able to collect the heart because the carcasses were removed from the cages. The remaining 9% of the mice had no evidence of coronary atherosclerosis and MI.

**Figure 7.**
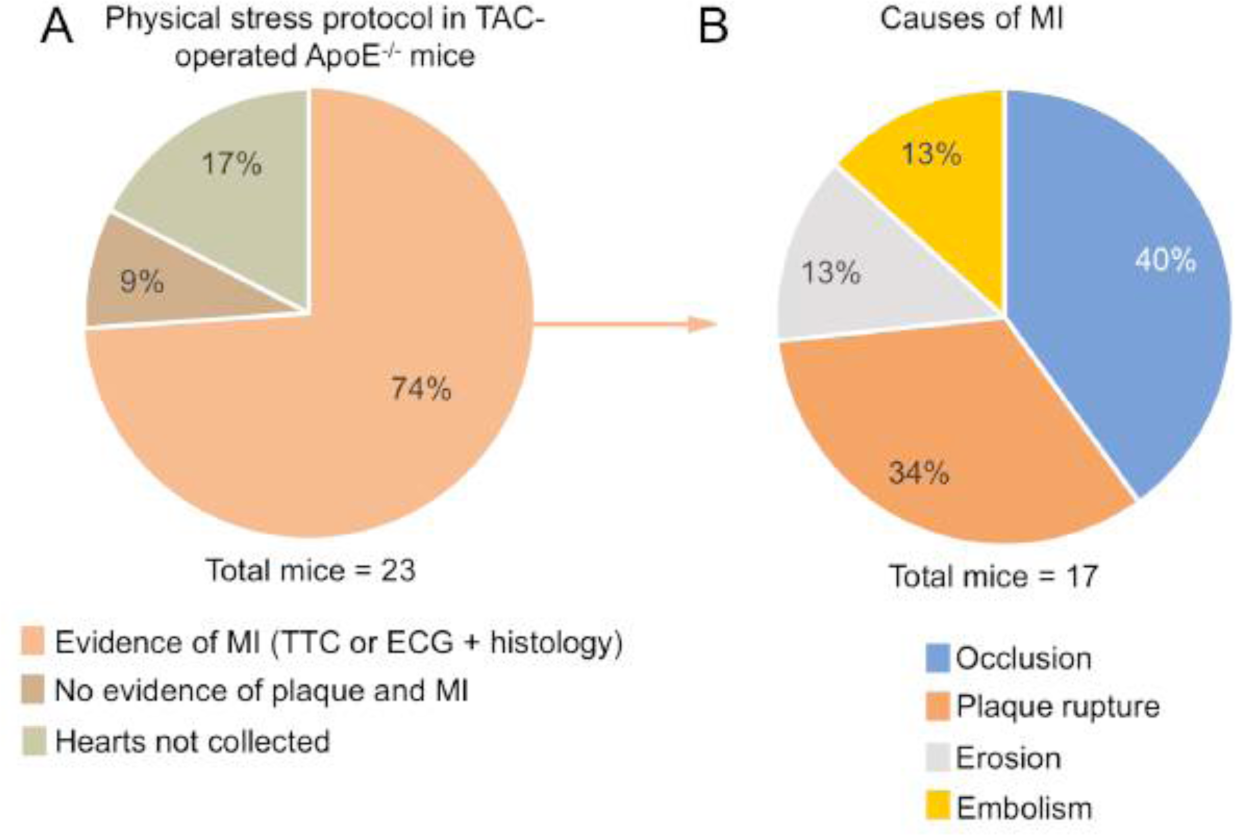
Causes of MI in TAC-operated ApoE^-/-^ mice subjected to physical stress. (A) Pie diagram of the TAC-operated ApoE^-/-^ mice died during the physical stress protocol: 74% of the mice presented evidence of MI determined by TTC staining or ECG and histological analysis; 9% did not show evidence of coronary plaque and MI, and 17% of the mice were discarded by RARC and therefore the heart were not collected. (n=23 TAC-operated ApoE^-/-^ mice). (B) Pie diagram illustrating the causes of MI of the 74% of animals dead during the physical stress: 40% of the mice showed coronary occlusion, 34% had evidence of plaque rupture; 13% of plaque erosion, 13% coronary embolism (n=17 TAC-operated ApoE^-/-^ mice).

We were able to define and quantify the causes of MI as follow: 47% of the mice presented clear evidence of coronary thrombosis, due to plaque rupture (34%) and plaque erosion (13%); 40% of the mice showed plaque occlusion with evidence of chronic myocardial fibrosis. Two mice had abrupt occlusion of coronary arteries by thrombus with no evidence of atherosclerosis, therefore classified as coronary embolism (13%).

## Discussion

For long time, establishing a mouse model of coronary atherosclerosis and its progression has been the “*holy grail”* in atherosclerosis research. ApoE^-/-^ and LDLR^-/-^ mice have been extensively used to study the pathogenesis of atherosclerosis(23) and provided great insights into the mechanisms of human disease. Studies in mice identified different genes involved in atherogenesis, and some of them have been confirmed in human pathology, including genes involved in triglyceride-rich lipoprotein metabolism(24, 25), and PCSK9 (protein convertase subtilisin/kexin type 9)(26, 27). Mouse models have proven to be very valuable to study disease mechanisms; however, many limitations exist that need to be taken into account, especially considering the challenges in translating the biology of atherosclerosis.

To develop atherosclerotic lesions, mice require cholesterol levels (∼2000 mg) that far exceed those commonly present in humans(9). More importantly, murine atherosclerosis develops in the aorta and its proximal branches, whereas coronary arteries are typically spared. The hemodynamics of the aorta and coronary arteries, as well as the origins of their SMC differ dramatically. SMC of coronary arteries originate from the proepicardial organ while the ones of descending aorta arise from somatic precursors(28, 29). Furthermore, plaque rupture, erosion and thrombus formation, which are the most critical complications of human atherosclerosis, are not present in current models (23, 30, 31). Whereas patients are treated at late stage of the disease when symptoms arise, mechanisms studied in mice focus on the development of atherosclerosis more than the progression of it. The difficulties in translating some of the studies in atherosclerosis has been ascribed, at least in part, to the aforementioned limitations of current mouse models(4, 9, 32, 33). To date, few attempts have been made to “model” coronary atherosclerosis and its sequelae in mice(10, 11), without realistic success.

Our study identifies the experimental conditions to induce the formation of coronary atherosclerosis and myocardial events, by combining moderated hypercholesterolemia (ApoE^-/-^, on chow diet) with increased afterload (TAC surgery), two major risk factors for CAD(34-37) and its complications. We have systematically characterized the topography, the magnitude and lipid composition of coronary plaques, in particular of the LAD (Fig. 1). The major novelty of our study is the formation of coronary lesions in mice exposed to specific pathological conditions (high cholesterol and pressure), which can progress over time and lead to MI under physical stress conditions (tread mill run), mainly due to coronary disruption and thrombosis, and coronary occlusion (Fig. 7). Physical exertion(38) and emotional stress may act as external triggers for acute MI(39). Similarly, in TAC-operated ApoE^-/-^ mice, physical stress induced myocardial events, in most of the mice (74%), during or within 48h from the treadmill run.

Most emphasis has been placed on the role of endothelium and macrophages in the initiation and progression of early atherosclerosis. However, human autopsy studies(40, 41), and *in vivo* data suggest an important role of SMC in the initial phase of atherosclerosis. As macrophage, SMC are known to express a variety of cholesterol uptake receptors, including LDL receptor(42), CD36(43), VLDL receptor(44), and therefore can form foam-like cells and participate to atherogenesis. SMC are also able to express adhesion molecules and retain inflammatory cells into the vascular wall. In early coronary lesions of ApoE^-/-^ mice at 8 week post-TAC, in addition to macrophages and EC, lipid accumulation were found in SMC of the media (Fig. 1), suggesting an active role of coronary SMC in the development of atherosclerosis in mice. Interestingly, as in human, the early coronary atherosclerosis was tightly associated with neointimal proliferation.

ApoE^-/-^ mice on high fat diet showed evidence of buried plaques in the innominate artery(45, 46), but no signs of the coronary lesions and disruption (47, 48), which is the cause of about 75% of coronary thrombi formation in humans(49, 50). Our study is the first demonstrating coronary plaque formation and progression towards disruption (rupture and/or erosion) and occlusion, which were accountable for 87% of the myocardial infarctions.

It is generally accepted the macrophages play an important role in the progression of atherosclerosis, by contributing as an important source of enzymes catabolizing collagen, a key component of the fibrous cap of the plaque(9, 51, 52). Interestingly, immunofluorescent staining (CD68 for macrophages, and α-SMA for SMC) evidenced a robust accumulation of macrophages into the ruptured plaque, and specifically adjacent to the points of break of the thin fibrous cap (Supplementary Fig. 2E). We are aware that these lineage markers for SMC (α-SMA) and macrophages (CD68) are promiscuous(53), as some cells are both, CD68 and α-SMA positive. The aim of these staining was not to quantitatively distinguish between these two populations, but to show the presence of macrophages in the ruptured plaques. Interestingly, in Suppl. Fig. 2, it is noticeable that the majority of CD68 and α-SMA staining do not co-localize, but only a minor population is double positive. Therefore, even considering these limitations, our data support the abundance of macrophages into the ruptured plaque, in correspondence to the thinning of the fibrous cap and breaking points.

In 40% of the mice studied, we found evidence of coronary ischemia based on ECG monitoring that revealed ST-elevation in some cases, and arrhythmic events such as accelerated idioventricular rhythm, in others. Indeed, we found stenosis and occlusions of the LAD arteries as well as other major coronary arteries (diagonal and marginal in our mouse model. It is possible that under physical stress, demand-related ischemia due to coronary artery disease and prior infarctions resulted in demise. Some mice also demonstrated acute MI potentially triggered by increased shear stress from the exercise. Prolonged ischemia has been shown to induce vasoconstriction in animals and human(54-56).

Lastly, in a small percentage of the infarcted hearts we found no evidence of coronary atherosclerosis with an abrupt interruption of blood flow by thrombus (Fig. 6). In human, coronary artery embolism occurs at low frequency and is commonly caused by atrial fibrillation(21, 57). Interestingly, ApoE^-/-^ mice following TAC, presented low rate of coronary embolism and we did not observe atrial fibrillation in our mice. We do not know the origin of the embolus but it is possible that the left ventricular dysfunction in our model contributed to formation of left ventricular thrombus.

In conclusion, the coexistence of high cholesterol and increased afterload can induce the formation of coronary lesions in ∼95% of ApoE^-/-^ mice, and leads to plaque disruption and occlusion in more than 70% of the mice (Figure 7). This pathological phenotype recapitulates major complications of human coronary atherosclerosis, specifically the formation in mice of coronary lesions that can progress to disruption, leading to thrombosis, or occlusion, ultimately causing myocardial events.

This study presents for the first time, a reliable tool to investigate not only the pathogenesis of coronary atherosclerosis, and its progression in mice, but also potential therapeutic targets.

## Methods

### Minimally invasive TAC

WT C57BL/6J and ApoE^-/-^ male mice (11-14 weeks of age; body weight 25-27 g) were subjected to TAC or sham operation (58). Briefly, mice were anesthetized using a single i.p. injection of ketamine (100 mg/kg) and xylazine (5 mg/kg). A topical depilatory was applied to the chest, and the area was cleaned with betadine and alcohol. A horizontal incision 0.5 to 0.8 cm in length was made at the second intercostal space. After retracting the thymus, the aortic arch was visualized with a dissecting scope (Zeiss Discovery.V8 Stereo) at low magnification. A 7-0 nylon ligature was tied between the innominate and left common carotid arteries with an overlying 28-gauge needle, which was then rapidly removed, leaving a discrete region of stenosis. In sham-operated animals the ligature was just tied loosely around the aorta.

### Cholesterol levels

Total cholesterol was measured in plasma from sham- or TAC-operated WT and ApoE^-/-^ mice, using a Cholesterol E kit (WAKO Life Science) according to the manufacturer’s instructions. Briefly, whole blood was collected in EDTA via cardiac puncture. Plasma was obtained by centrifugation at 1,000 × g for 15 min at 4 °C.

### Stress-induced MI protocol

Starting from 8-week post-TAC, ApoE^-/-^ mice were placed on the treadmill (Columbus Instrument) to run for 4 min/day, once a week for 12 weeks at a maximal sustained running speed of 13-14 m/min with 15° of inclination. Because this test is a form of forced exercise, when sometimes the mice stopped for short periods, they were gently tapped or lifted to encourage resumption of running. A minimal electric shock was also applied in order to encourage the run on the treadmill.

### Radiotelemetry for electrocardiographic analysis

Electrocardiograms (ECG) in conscious mice were measured using telemetry transmitters (ETA-F10 transmitter; Data Sciences International, St. Paul, MN). Briefly, the transmitter device, which communicates with the receiver wirelessly, was inserted subcutaneously through a dorsal incision in the abdomen of the animal, and the two ECG electrodes were placed subcutaneously along the superior right pectoral region (negative lead) and the left caudal rib position (positive lead) to approximate lead II of the Einthoven surface electrocardiogram. Mice were housed individually in a temperature - and humidity - controlled facility and maintained under a 12-hour dark and 12-hour light cycle and they were allowed to recover for 7 days after surgery. ECG traces were continuously recorded at baseline, during the stress-induced MI protocol and during the following 48 hours, using LabChart software (AD Instruments).

### Echocardiographic analysis

Cardiac dimensions and function were analyzed by transthoracic echocardiography using a Vevo 770 Imaging System (VisualSonics) as previously described(58). Mice were lightly anesthetized with inhaled isoflurane (0.2% in O_2_). Left ventricle M-mode was used, and all measurements were obtained from 3 to 6 consecutive cardiac cycles, and the average values were used for analysis. Left ventricle end-diastolic (LVDd) and end-systolic (LVDs) dimensions were measured from the M-mode traces, and fractional shortening (FS) was calculated as follows: [(LVDd − LVDs)/LVDd]. Diastolic measurements were taken at the point of maximum cavity dimension, and systolic measurements were made at the point of minimum cavity dimension, using the leading-edge method of the American Society of Echocardiography(59).

### Oil red-O staining

To assess lipid deposition in the coronary arteries, frozen heart sections were stained with Oil Red O (Sigma-Aldrich) as previously described(60). Briefly, heart cryosections were washed in PBS, stained with 0.2% Oil Red O in 60% isopropanol for 20 min, and washed with 60% isopropanol. After counterstaining, performed with Mayer’s hematoxylin, sections were washed and coverslipped with aqueous mounting medium. Images were acquired by using a Zeiss Axio Observer.Z1 microscope and the area of Oil red-O staining was calculated by using ImagePro (Media Cybernetics). Coronary stenosis were calculated by measuring the area of the lumen (A_L_), the internal elastic lamina (A_IEL_) and calculate the percent of stenosis as follows: (A_IEL_-A_L_)/A_IEL_ x 100. Oil Red-O percent in the plaque stenosis was calculated as the ratio of positive Oil Red-O staining area to the plaque area (%).

### Masson trichrome staining

Myocardial fibrotic tissue was determined in TAC-operated WT and ApoE^-/-^ mice, using a Masson’s trichrome stain ki (Polysciences, Inc.) according to the manufacturer’s instructions.

### Histology and immunostaining

Mouse hearts were collected no more than 1h post-mortem, without perfusion and fixed with 4% paraformaldehyde (PFA) overnight at 4°C. The following day, PFA 4% was replaced by sucrose 30% and left overnight at 4°C on rotation. The hearts were divided into 3 parts (base, center, and apex), immerged overnight with OCT: sucrose 30% (1:1 ratio), and consequently embedded in OCT. The whole heart was sectioned, with sets of 10 consecutive sections of 10 µm collected every 30 µm. Heart sections every 260 µm were routinely stained with H&E, and coverslipped with aqueous mounting medium.

For immunofluorescence analysis, the heart sections were permeabilized in 0.5% Triton X-100, in a solution of 5% BSA 5%, followed by blocking in BSA 5% (45 min at room temperature). Next, the heart sections were incubated overnight at 4°C with anti-CD41 antibody (1:100, 553897; BD Pharmingen), anti-alpha smooth muscle actin antibody (1:200, 180186; Life technologies), anti-CD31 (1:100, 553370; BD Biosciences), anti-CD68 antibody (1:200, MCA1957; BioRAD) and anti-collagen I antibody (1:200, ab43710; Abcam) in BSA 5%. After 3 washes in PBS, secondary antibodies Alexa Fluor 568-conjugated donkey anti-mouse IgG (1:200, ab34710; Abcam), Alexa Fluor 647-conjugated goat anti-rat IgG (1:500, A21247, ThermoFisher Scientific) Cy5-labeled donkey anti-rabbit IgG (1:500, 711-175-152; Jackson ImmunoResearch) were added for 1h at room temperature. Nuclei were counterstained with Hoechst 33258 (1:10000, 382061; Calbiochem). Immunofluorescence images of heart sections were captured in a Z-stack with 0.48-µm steps using a Zeiss Axio Observer.Z1 microscope followed by deconvolution using the ZEN pro 2012 software (Carl Zeiss).

### Study Approval

All of the studies were performed according to protocols approved by the Weill Cornell Institutional Animal Care and Use Committee.

## Statistics

Statistical analysis was performed with GraphPad Prism 7.0 software (San Diego, CA, USA). All data are reported as mean ± SEM; Statistical analysis was performed with one-way ANOVA followed by Dunnett’s post-hoc test when comparing more than two groups of data (Fig 1H and Fig. 2G). Data were considered statistically significant when a value of at least p<0.05 was achieved.

## AUTHOR CONTRIBUTIONS

ADL, AM and YZ designed the studies. AM, YZ, LR and JI conducted experiments and acquired data. ADL, AM, YZ and JI analyzed data. ADL and AM wrote the manuscript. JI contributed to edit the manuscript.

## ACKNOWLEDGMENTS

This work was supported by National Institute of Health grant R01HL126913, Harold S. Geneen Charitable Trust Awards Program for Coronary Heart Disease research to A.D.L. We are very thankful to Dr. Domenick J. Falcone and Dr. Roberto Levi for critical input and scientific discussion of the manuscript.

## Abbreviations

ApoE^-/-^: Apolipoprotein E deficient
AV: Aortic valve
CAD: Coronary artery disease
ECG: Electrocardiogram
FS: Fractional shortening
LDLR^-/-^: Low density lipoprotein receptor deficient mice
LAD: Left anterior descending artery
MI: Myocardial infarction
TAC: Transverse aortic constriction

