## Supplemental Figures for "Pressure overload leads to coronary plaque formation, progression and myocardial events in ApoE^-/-^ mice"

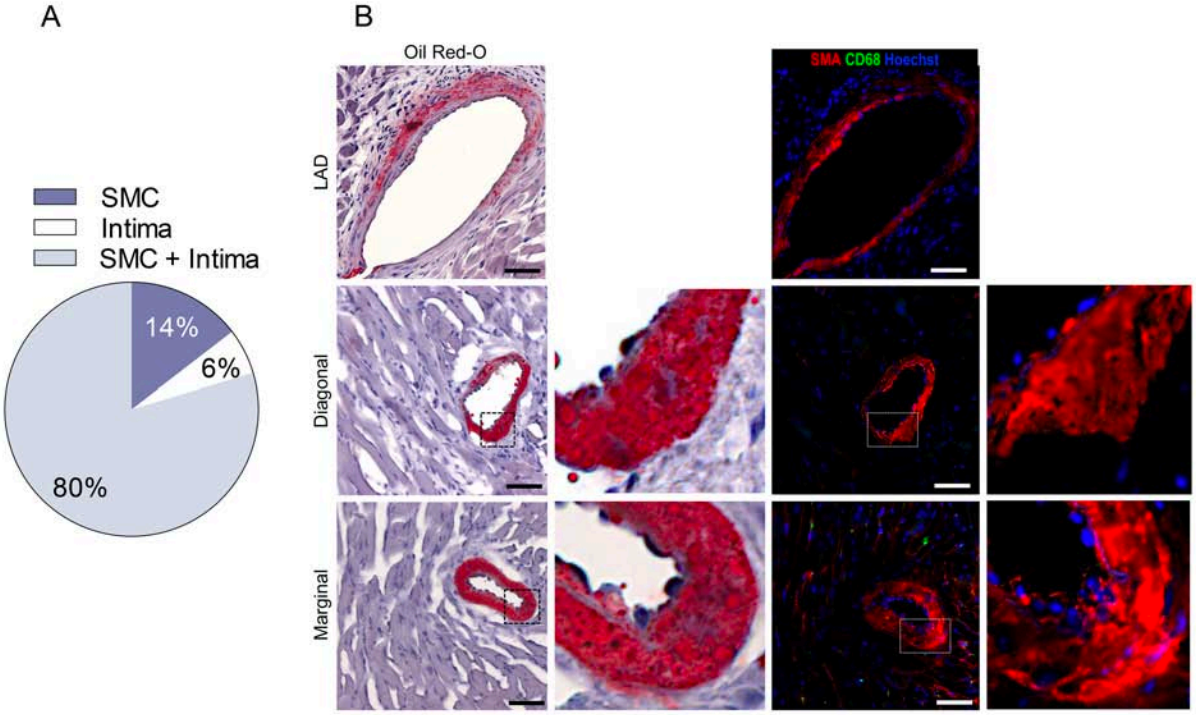


**Supplemental Figure 1. Lipid deposition in coronary early lesions of ApoE$^{-/-}$ mice at 8-week post-TAC .** (A) Pie chart representing the distribution of lipids accumulation into the coronary wall: in 80% of early lesions lipids accumulate in both SMC and intima; in 14% of the early lesions lipids accumulate in SMC; and in 6% of the cases in the intima (n=103 coronary lesions from 10 mice). (B) Myocardial sections of ApoE$^{-/-}$ mouse at 8-week post-TAC were stained with oil red-O & hematoxylin, and immunofluorescently stained with $\alpha$-SMA (SMC), CD68 (monocyte/macrophage) and Hoechst (nuclei). Oil red-O showed lipid accumulation in the media of proximal LAD, and diagonal and marginal arteries. Immunofluorescent staining for $\alpha$-SMA (red) confirms the accumulation of lipids into smooth muscle cells, while co-staining for CD68 is negative. Nuclei were stained with Hoechst (blue). Scale bars: 50 μm.

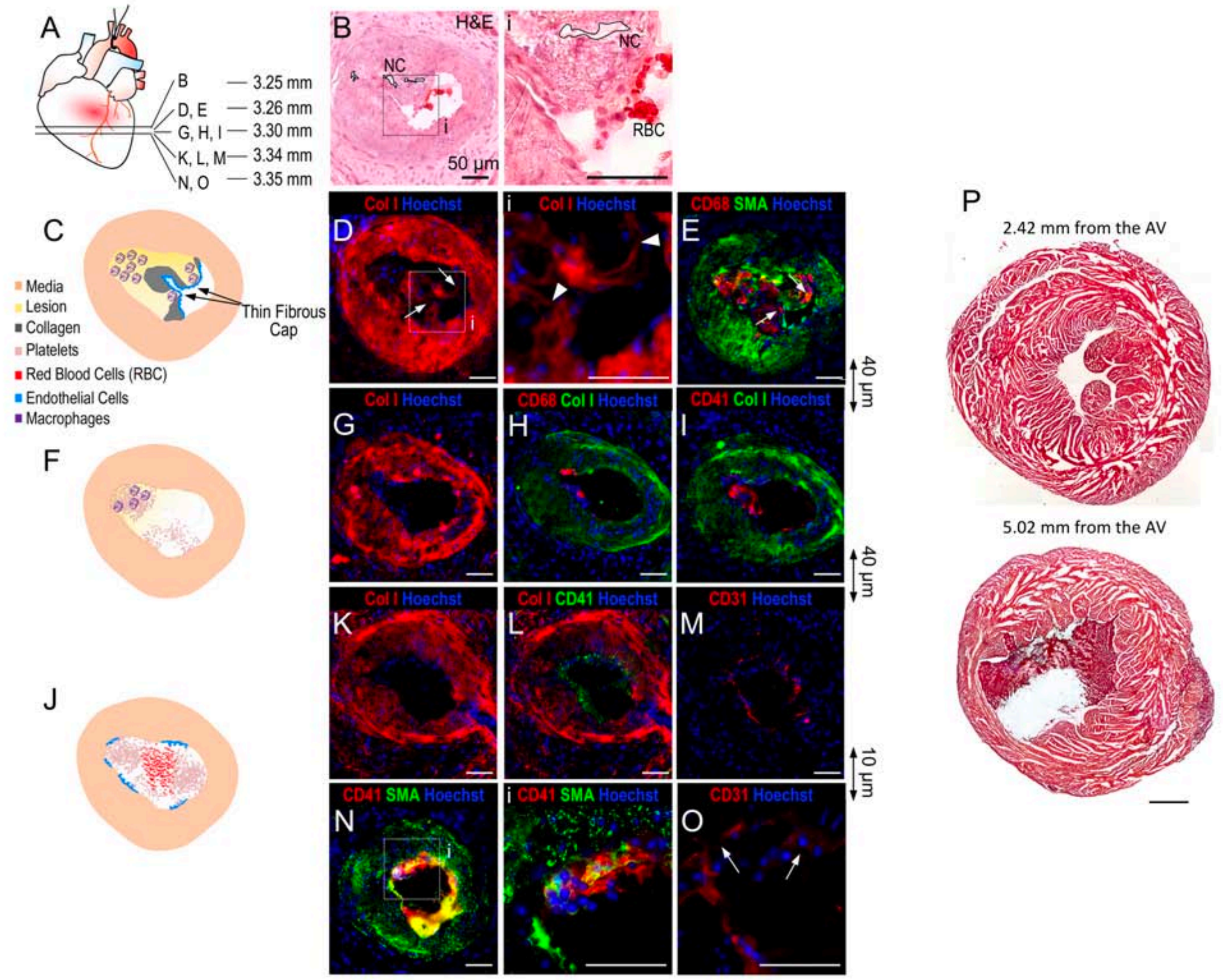


**Supplemental Figure 2. Coronary plaque rupture and thrombotic occlusion in ApoE$^{-/-}$ mice post-TAC.** (A) Scheme of the localization of immunofluorescently labeled heart sections proximal and distal to the section with thrombus shown in panel B and inset i. (B) At 3.25 mm from the AV there is a major lesion as shown by the H&E staining and inset i. (D,E) IF staining showed a very thin fibrous cap (Col I and $\alpha$-SMA staining) in two points (panel D, inset i, arrowheads). (E) IF staining for CD68 shows macrophage accumulation in close proximity to the thin fibrous cap (arrows). (C) Schematic representation of the plaque in the LAD based on panels D,E. (G-I) IF staining of consecutive myocardial sections for Col I (red), macrophages (CD68, red), platelets (CD41, red) and smooth muscle cells ($\alpha$-SMA, green) reveal that the major part of plaque detached from the coronary wall. In panel H is still possible to notice some remaining CD68 positive cells of the lipid core of the plaque, and in panel I platelets aggregates interacting with CD68-positive cells and ruptured plaque. (F) Schematic representation of the IF staining in panels G-I. (K-O) Consecutive downstream sections showing platelet aggregates in intimate contact with the plaque downstream the point of rupture. (O) It is possible to observe the disruption of the endothelial layer, possibly by erosion, and platelets aggregates in correspondence of the disrupted endothelium in contact with SMC (panel N, inset i). (J) Schematic cartoon representing the IF staining in panels K-O of the thrombotic event in the LAD. Scale bars: 50 μm. (P) Masson's trichrome staining of the medium and apex of the heart at the indicated distance from the AV.
